## Supplementary Material for "3D-MINFLUX nanoscopy reveals distinct allosteric mechanisms for activation and modulation of PIEZO1 by Yoda1"

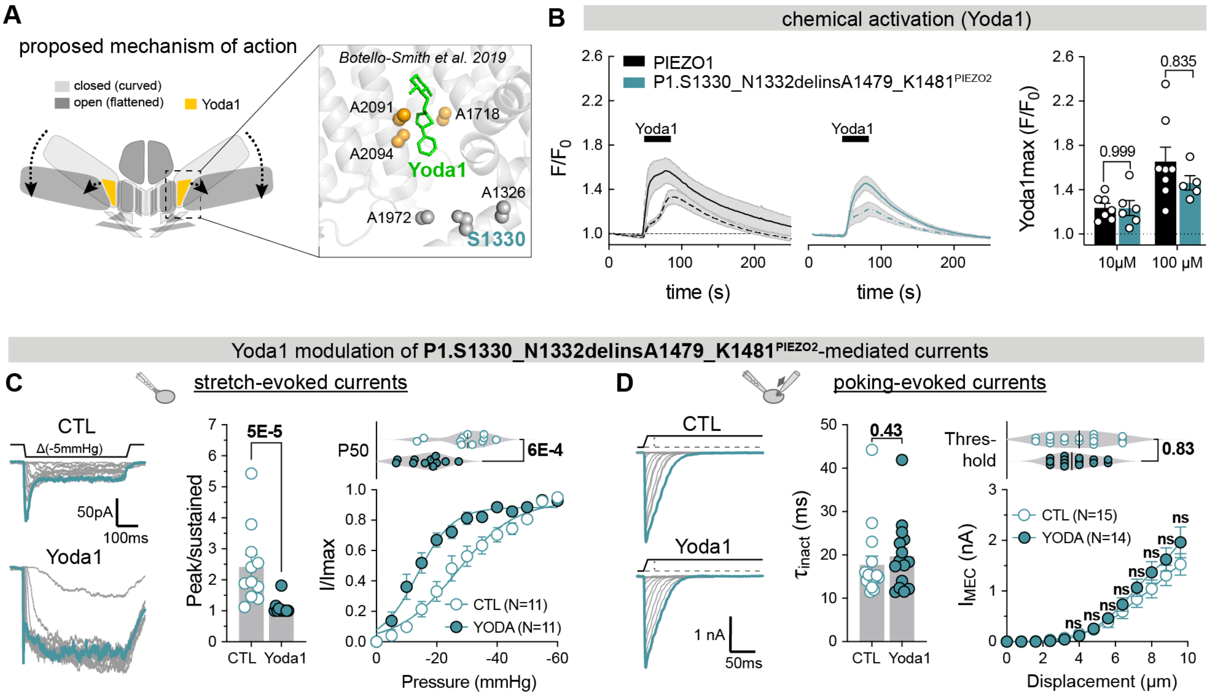


**Supplementary Figure 1 | Mutation of a possible Yoda1 binding site at the beam–THU8/9 interface does not alter Yoda1 sensitivity (A)** Cartoon depicting the proposed mechanism of action of Yoda1 on PIEZO1 (left) and close-up view of the THU8–9 interface with previously proposed residues lining putative Yoda1 binding sites highlighted in sphere representation. **(B)**, time course of $Ca^{2+}$-influx (F/F0) evoked by 10 and 100 µM Yoda1 in cells expressing PIEZO1 (left) and P1.S1330_N1332delinsA1479_K1481$^{PIEZO2}$ (center) assessed by GCamp8 imaging together with comparison of maximum responses (right). **(C)** Modulation of P1.S1330_N1332delinsA1479_K1481$^{PIEZO2}$ stretch-evoked currents by 30µM Yoda1. Example traces evoked by incrementing pressure stimuli (left), comparison of peak/sustained ratio (middle) using Mann-Whitney test (CTL = 2.4 ± 1.2, N=11 vs Yoda1 = 1.1 ± 0.2, N=11, P=0.00005), pressure-response curves (i.e. peak current amplitude at indicated pressure normalized to maximal response, bottom right) and comparison of $P_{50}$ values in the absence and presence of Yoda1 using Student's t-test test (CTL = -29.3 ± 8.2 mmHg, N=11 vs Yoda1 = -16.5 ± 6.5 mmHg, N=11, P=0.0006). **(D)** Modulation of P1.S1330_N1332delinsA1479_K1481$^{PIEZO2}$ poking-evoked currents in whole-cell recordings by 30µM Yoda1. Example traces evoked by incrementing (Δ 800nm, left), comparison of inactivation time constants obtained with exponential decay fit (middle) using XY test (CTL = 17.7 ± 8.2 ms, N=16 vs Yoda1 = 19.64 ± 7.9 ms, N=15, P=0.429), displacement-response curves (i.e. peak current amplitude vs. indicated stimulus magnitude; bottom, right) and comparison of mechanical activation thresholds using Mann-Whitney test (CTL = 3.95 ± 1.6 µm, N=16 vs Yoda1 = 3.77 ± 1.1, N=14, P=0.825).

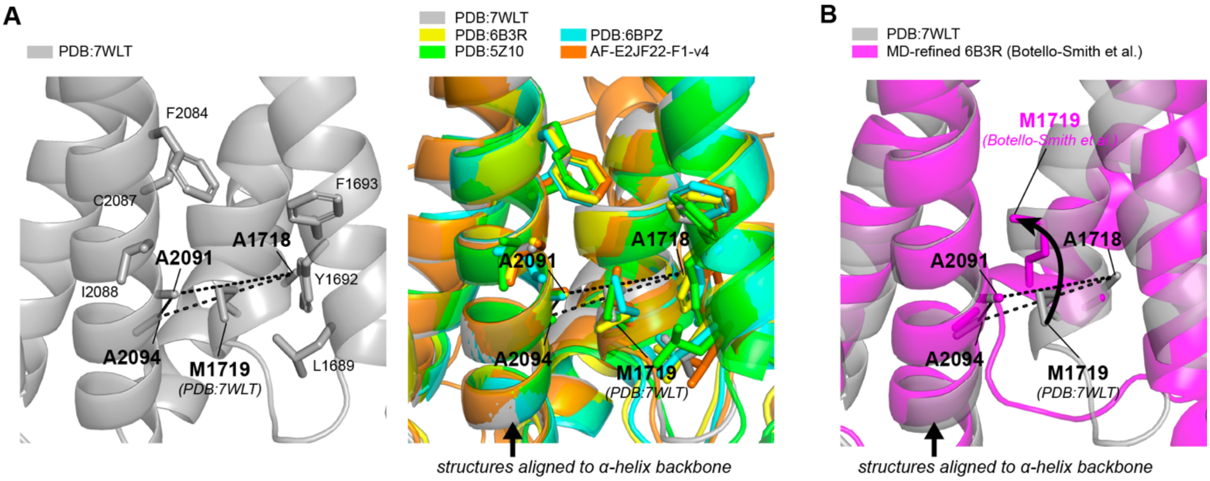


**Supplementary Figure 2 | Comparison of THU8–9 interface in available PIEZO1 structures. (A)** left, close-up side view of the THU8–9 interface in PDB:7WLT with residues A2091, A2094 and A1718, supposedly lining the Yoda1 curved-state binding site highlighted in stick representation. Note, the space between the three residues is occupied by the side chain of M1719. Right, overlay of all previously resolved PIEZO1 cryo-EM structures (7WLT, 6B3R, 5Z10, 6BPZ) as well as the Alphafold prediction (AF-E2JF22-F1-v4), demonstrating the same structural arrangement of side chains in all five structures. Note, for better comparison the structures were aligned to the indicated helix of the 7WLT structure. **(B)**, Overlay of the 7WLT structure used for binding pocket and docking pose analysis here (7WLT, grey) and the molecular dynamics (MD)simulation-refined structure previously analysed by Botello-Smith and colleagues. Note, there is a subtle yet relevant difference in the orientation of the M1719 side chain and the α-helix in which it resides, such that the space between A2091, A2094 and A1718 is vacant in the MD-refine structure allowing Yoda binding.

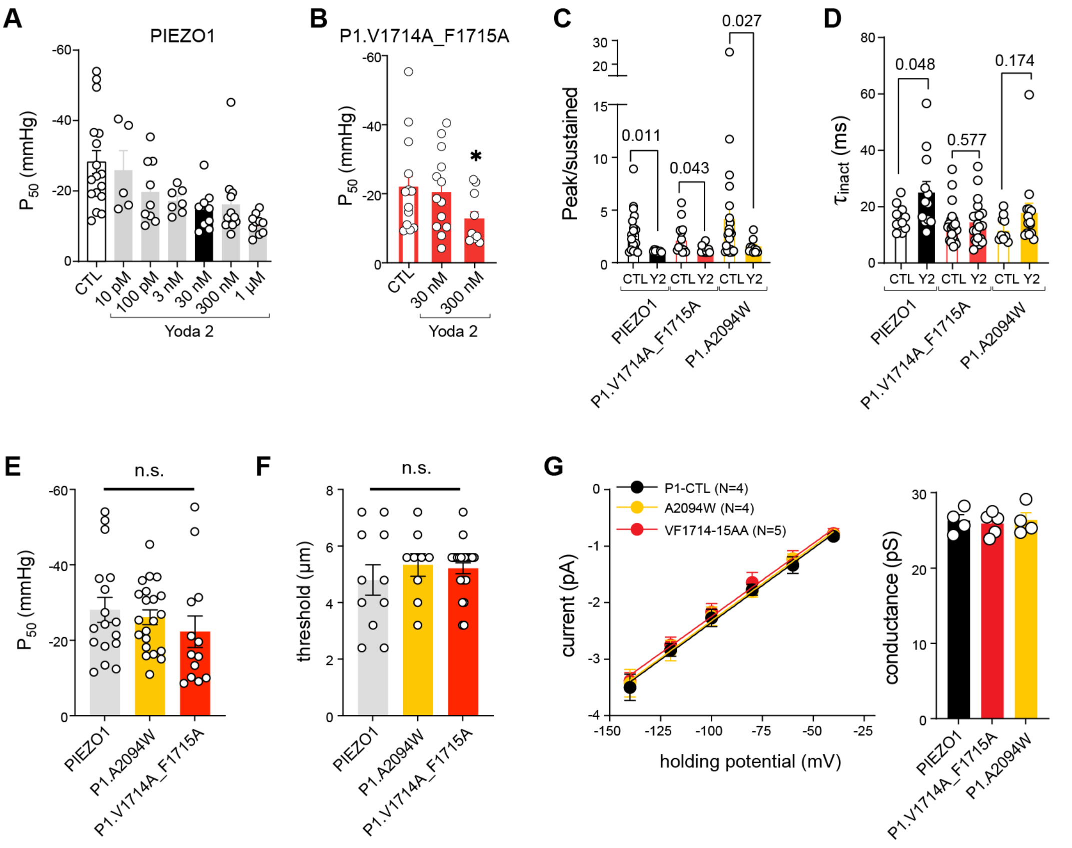


**Supplementary Figure 3 | Electrophysiological characterization of Yoda2 modulatory effect on PIEZO1 and associated mutants. (A)** Comparison of the mean ± s.e.m. P50 for PIEZO1 Control (N=17) and Yoda2 at various concentration (N=5-12), determined from stretch-evoked currents in N2a-P1KO cells. **(B)** Comparison of the mean ± s.e.m. P50 for PIEZO1.V1714A_F1715A Control (N=13) and Yoda2 at 300nM (N=9), Mann-Witney test CTL vs 300nM (P=0.0433). **(C)** Kinetic analysis of stretch-evoked currents. Comparison of the mean ± s.e.m. Peak/sustained ratio for PIEZO1 and associated mutants in the presence or absence of 30nM Yoda2, Mann-Witney test. **(D)** Kinetic analysis of whole-cell poking-evoked currents. Comparison of the mean ± s.e.m. inactivation time constant for PIEZO1 and associated mutants in the presence or absence of 30nM Yoda2, unpaired t-test. **(E)** Comparison of the control (untreated) P50 from stretch-evoked currents of PIEZO1, A2094W and V1714A_F1715A, one-way ANOVA (P=04429. **(F)** Comparison of the control (untreated) threshold of whole-cell poking-evoked currents of PIEZO1, A2094W and V1714A_F1715A, one-way ANOVA (P=0.5769). **(G)** Current/voltage relationship of single-channel amplitude of PIEZO1, A2094W and V1714A_F1715A, fitted with a linear regression (left). Comparison of the mean ± s.e.m. single channel conductance (right), Kruskall-Wallis test (P=0.9523).

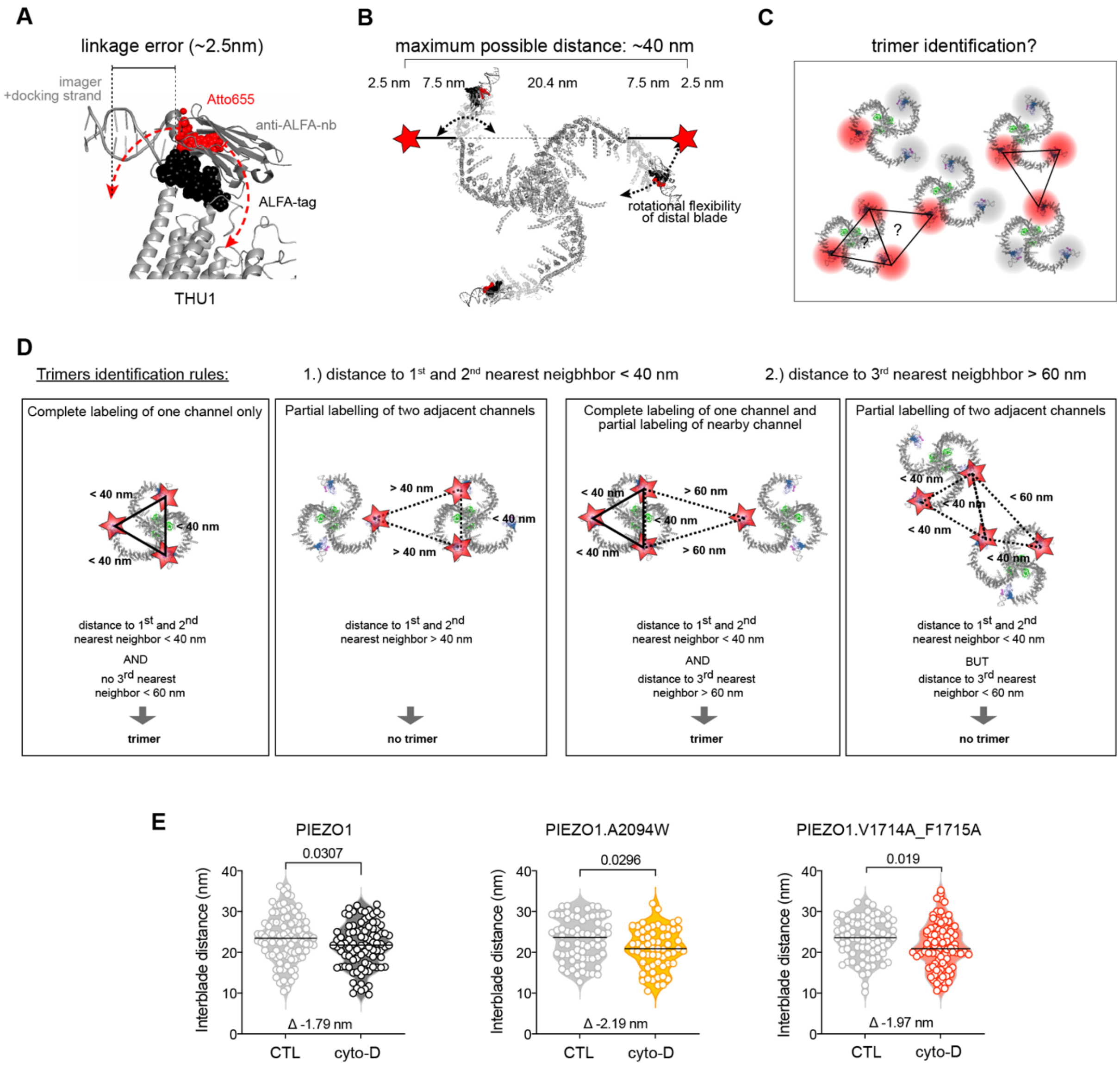


**Supplementary Figure 4 | PIEZO trimer identification with 3D-MINFLUX. (A**) Cartoon illustrating the labelling strategy and its associated linkage error. The position of the extracellular ALFA tag is depicted, with its associated nanobody with a DNA strand and the complementary DNA strand with Atto655 fluorophore. **(B)** Representation of a full-length PIEZO1 (top view) and the possible maximum interblade distance measured experimentally. **(C)** Schematic representation of the trimer identification problem due to partial labelling and proximity of neighbouring PIEZO1 channels. **(D)** Trimer identification pipeline. **(E)** Comparison of the mean ± s.e.m. interblade distance of the identified trimers in untreated cells or after addition of cytochalasin-D for PIEZO1 (N=98 and 93), A2094W (N=70 and 61) and V1714A_1715A (N=78 and 87), with unpaired t-test or Mann-Witney test (A2094W).

| Primers | 5' --> 3' | Template | Generated construct |
| --- | --- | --- | --- |
| P1-A2094W-fw | atttactttgccctgtcctggtaccagatccgctgtggc | PIEZO1-mScarlet | PIEZO1-A2094W-mScarlet |
| P1-A2094W-rv | gccacagcggatctggtaccaggacagggcaaagtaaat | | |
| P1-A2094W-fw | atttactttgccctgtcctggtaccagatccgctgtggc | PIEZO1-ALFA-mGreenLantern | PIEZO1-ALFA-A2094W-mGL |
| P1-A2094W-rv | gccacagcggatctggtaccaggacagggcaaagtaaat | | |
| P1-KSIN-P2-fw | aatgcagccaacctgaaggcggtcaaGttccatcgccagattgag | PIEZO1-mScarlet | PIEZO1-S1330-N1332del-insA1479-K1481-P2 |
| P1-KSIN-P2-rv | ctcaatctggcgatggaaCttgatcgccttcaggttggctgcatt | | |
| P1-VF1714-15AA-fw | gtgctgcccgtgcttgCgGCcctgtgggccatgctg | PIEZO1-mScarlet | PIEZO1-VF1714-15AA-mScarlet |
| P1-VF1714-15AA-rv | cagcatggcccacaggGCcGcaagcacgggcagcac | | |
| P1-VF1714-15AA-fw | gtgctgcccgtgcttgCgGCcctgtgggccatgctg | PIEZO1-ALFA-mGreenLantern | PIEZO1-ALFA-VF1714-15AA-mGL |
| P1-VF1714-15AA-rv | cagcatggcccacaggGCcGcaagcacgggcagcac | | |
| P1-VF1714-15AA-fw | gtgctgcccgtgcttgCgGCcctgtgggccatgctg | PIEZO1-A2094W-mScarlet | PIEZO1-VF1714-15AA-A2094W-mScarlet |
| P1-VF1714-15AA-rv | cagcatggcccacaggGCcGcaagcacgggcagcac | | |

### Supplementary Table 1, DNA-primers used for cloning

| Iteration | Modality | Pattern diameter (nm) | photon limit | back-ground limit | dwell time (ms) | pattern repeat | Stickiness | CFR limit | laser power factor |
| --- | --- | --- | --- | --- | --- | --- | --- | --- | --- |
| 0 | hexagonal | 251 | 160 | 15000 | 1 | 1 | - | none | 1 |
| 1 | zline | 251 | 400 | 15000 | 1 | 1 | 2 | none | 1 |
| 2 | square | 251 | 100 | 10000 | 1 | 5 | 2 | none | 1 |
| 3 | zline2 | 251 | 50 | 10000 | 1 | 5 | 2 | none | 1 |
| 4 | square | 132 | 67 | 10000 | 1 | 5 | 2 | 0.9 | 2 |
| 5 | zline2 | 132 | 33 | 10000 | 1 | 5 | 2 | none | 2 |
| 6 | square | 66 | 67 | 10000 | 1 | 5 | 2 | 0.8 | 4 |
| 7 | zline2 | 66 | 33 | 10000 | 1 | 5 | 2 | none | 4 |
| 8 | square | 35 | 100 | 10000 | 1 | 5 | 2 | none | 6 |
| 9 | zline2 | 35 | 50 | 10000 | 1 | 5 | 2 | none | 6 |

### Supplementary Table 2, 3D-MINFLUX scan parameters
